# Integrating multiplexing into confineable gene drives effectively overrides resistance in *Anopheles stephensi*

**DOI:** 10.1101/2025.08.15.670499

**Authors:** Mireia Larrosa Godall, Lewis Shackleford, Matthew P. Edgington, Philip T. Leftwich, James C. Y. Luk, Joshua Southworth, Stewart Rosell, Jake Creasey, Jack Aked, Katherine Nevard, Alexander Dodds, Morgan Mckee, Eunice Adedeji, Estela Gonzalez, Joshua X. D. Ang, Michelle A. E. Anderson, Luke Alphey

## Abstract

*Anopheles stephensi* is a major malaria vector mainly present in southern Asia and the Arabian Peninsula. Since 2012 it has invaded several countries of eastern Africa, stimulating urgent efforts to develop more efficient strategies for vector control such as CRISPR/Cas9-based homing gene drives. Target site resistance is a significant challenge to the deployment of these systems. When a double-stranded break is repaired by NHEJ, it can lead to mutations which destroy the target site, making that allele unrecognizable to the sgRNA and resistant to further cleavage. The use of multiple sgRNAs has the potential to solve this issue. We performed experimental crosses to assess the homing and cutting efficiency of two different multiplexing strategies targeting the *cardinal* locus, in the presence and absence of a resistance allele. We found pre-existing mutations at one sgRNA target site did not significantly reduce the homing efficiency for either strategy. Modelling indicates that while both strategies can overcome resistance allele formation, the fitness of the drive-carrying alleles is a critical factor in determining the overall performance and persistence of a split drive.

## Introduction

In 2021 there were approximately 247 million cases and 619,000 deaths due to malaria, of which 95% and 96%, respectively, occurred in Africa^1^. The primary vectors of this disease in the African continent are *Anopheles gambiae s*.*s*., *An. coluzzii, An. arabiensis* and *An. funestus*; all of which have life cycles adapted to rural settings, resulting in malaria being an overwhelmingly rural disease^2^. In contrast, *An. stephensi*, the main malaria vector in India and Pakistan, is better adapted to urban environments. Its unique ability to locate clean water, usually in storage tanks, to lay its eggs^2^ instead of more typical larval breeding sites used by African native vectors such as puddles and ditches, allows it to avoid polluted water (e.g. oil and sewage) in urban areas^3^. This adaptation allowed *An. stephensi* to thrive in urban areas, driving malaria outbreaks in cities across India, Pakistan, the Arabian Peninsula and Iran^4^. Since the first reported finding in Djibouti in 2012, *An. stephensi* has invaded west through Africa, expanding its range into Ethiopia and Sudan in 2016, Somalia in 2019, Nigeria in 2020, Kenya and Ghana in 2022, and recently in the Republic of Niger (2025)^5–9^. This threatens the 40% of sub-Saharan Africans living in urban areas and puts an additional 126 million people at risk^10^. Vector control interventions such as indoor residual spraying (IRS) and insecticide-treated nets (ITNs) are currently used to control malaria transmission. However, the efficiency of these strategies has been threatened by an increase in insecticide resistance by mosquitoes^1,7,11^. As a result in 2023 the WHO released an initiative to stop the spread of this species in Africa, calling for prioritising research to evaluate new tools to control *An. stephensi*^11^.

The ability to modify DNA using recent available tools such as clustered regulatory interspaced short palindromic repeats-associated protein 9 (CRISPR/Cas9) has given rise to alternative approaches that use genetic manipulation to reduce vector competence or suppress wild mosquito populations with relatively small initial releases^12,13^. The Cas9 endonuclease targets a specific sequence determined by the single guide RNA (sgRNA) and initiates a double-stranded break. The DNA is repaired by cellular mechanisms such as homology-directed repair (HDR) or non-homologous end joining (NHEJ). By using CRISPR/Cas9 to bias the inheritance of a cargo gene, populations of mosquitoes could be modified to be refractory to malaria infection^14^ or, by targeting a gene vital for fertility, they could suppress populations^15^. For both strategies, repair through the NHEJ pathway will often result in a change to the sequence of the target site, generating an allele which is resistant to further sgRNA recognition and cleavage by Cas9. Consequently, individuals which inherit these mutations are also immune to future drive allele conversion^16–20^. Such formation of cut-resistant alleles is more likely to occur when embryos contain maternally deposited Cas9 which may produce cuts early in development^16^. As a result, high rates of cut-resistant allele formation can threaten the success of any form of non-sex specific gene-drive, especially if the cargo gene carries a fitness cost such that the resistant allele could outcompete the drive allele in the population^16,21,22^.

One promising strategy to mitigate the threat of resistance alleles is to multiplex, e.g. designing a drive construct which expresses multiple sgRNAs targeting the same gene^20,23–27^. In the classical multiplexing strategy, a single construct expresses multiple sgRNAs which cut adjacent target sites and resistance allele formation at one or even multiple target sites would not halt the spread of a gene drive into the population as long as there is at least one remaining intact target site. This dramatically increases the number of NHEJ events required to form a fully drive resistant allele. This also decreases the probability and frequency of cut-resistant individuals generated in a population. These cut-resistant mutants can be functional (r1) or non-functional (r2) and pose a threat to a gene drive’s ability to spread and persist in a population^28^. One concern when utilizing these strategies is the effect of sequence heterology or dissimilarity between the homology arms of the donor template and the cut chromosome on drive efficiency, due to the requirement to resect the cut allele some distance to reach homologous sequences^26,27,29^. Multiplexing with multiple sgRNAs can also reduce the likelihood of mutations being functional as each target site is cut and repaired. Another significant drawback to the classical multiplexing strategy is that the entire target region between the two furthermost target sites might be deleted either by NHEJ or microhomology-mediated end joining (MMEJ) if two targets are cut simultaneously, removing all target sites in one step^20,22,30–32^. Therefore, alternative multiplexing strategies that allow multiplexing at the population level instead of the individual level have been proposed^23^. These strategies focus on targeting genes through multiple, distinct constructs as opposed to multiple adjacent sites targeted by a single construct. One of the proposed strategies, the additive strategy, is characterised by targeting different sites spaced so they are unlikely to segregate from each other through recombination but also are distant enough that homing at each target site can occur independently. Modelling of the additive strategy has shown that its use increases the number of generations in which gene drives persist at high frequencies compared to the classical multiplexing strategy^23^.

Genes in the ommochrome pathway have often been used as targets for homing-based gene drives in Anopheline mosquitoes. Genetic manipulation of these genes affects eye colour, making them experimentally tractable; null mutations of their homologues in *Drosophila melanogaster* are viable and fertile. These genes are *kynurenine 3-monooxygenase* (*kmo*), which resulted in a reduction in the survival after blood-feeding in homozygous *An. stephensi females*^16,33^, and *cardinal* (*cd*), which did not show any apparent reproductive or survival fitness cost in *An. gambiae mosquitoes*^14^. Based on this information *cd* was selected as a target gene for this study in *An. stephensi*. Here, we investigate multiplexing strategies to mitigate the risk of resistance allele formation targeting the *cd* gene. We demonstrate the ability of two strategies to continue homing in the presence of a resistance allele. However, we found that, likely due to extremely high cleavage rates, in the rare instances when homing did not occur, the classical multiplexing design frequently generated large deletions which removed several sgRNA target sites simultaneously. Modelling indicates that this type of deletion is the most frequent cause for the classical multiplexing design to fail^23^. We demonstrate through modelling that the additive strategy can successfully bypass this issue. However, this strategy is sensitive to the fitness costs of successive insertions.

## Results

### The classical multiplexing strategy gives high homing rates despite the requirement for resection

We have previously shown >98% inheritance using a transgenic line inserted in the 384 sgRNA target site expressing a single sgRNA from the As7SK promoter (*cd*^7SK^, here referred to as *cd*^*g*384^ to distinguish from other cd targets) and a line expressing Cas9 from the endogenous zpg locus (*zpg*^*3’Cas9*^)^34^. To analyse the feasibility of the classic multiplexing strategy to mitigate target site resistance, a construct carrying a *Hr5/IE1-ZsGreen*-*K10* fluorescent marker and four sgRNAs expressed by four different RNA pol III promoters^34^, with four adjacent targets in cd, was designed for HDR insertion into this same region of the cd gene. The target sites are within 136 nucleotides (nt), aiming to preserve homing efficiency while reducing the possibility of deletions that may remove the entire intervening sequence. The resulting plasmid was used to generate the *cd*^*g*338-384^ line through CRISPR/Cas9-mediated integration into cd (Figure 1A). A second construct and transgenic line containing the same homology arms (keeping the 136nt deletion of the additional sgRNA target sites) and marker cassette as the *cd*^*g*338-384^ line but only expressing one sgRNA recognizing the 384 cut site was also generated, *cd*^*g*384_del^ (Figure 1A). This line serves to determine the effect of resection on drive conversion efficiency.

**Figure 1.**
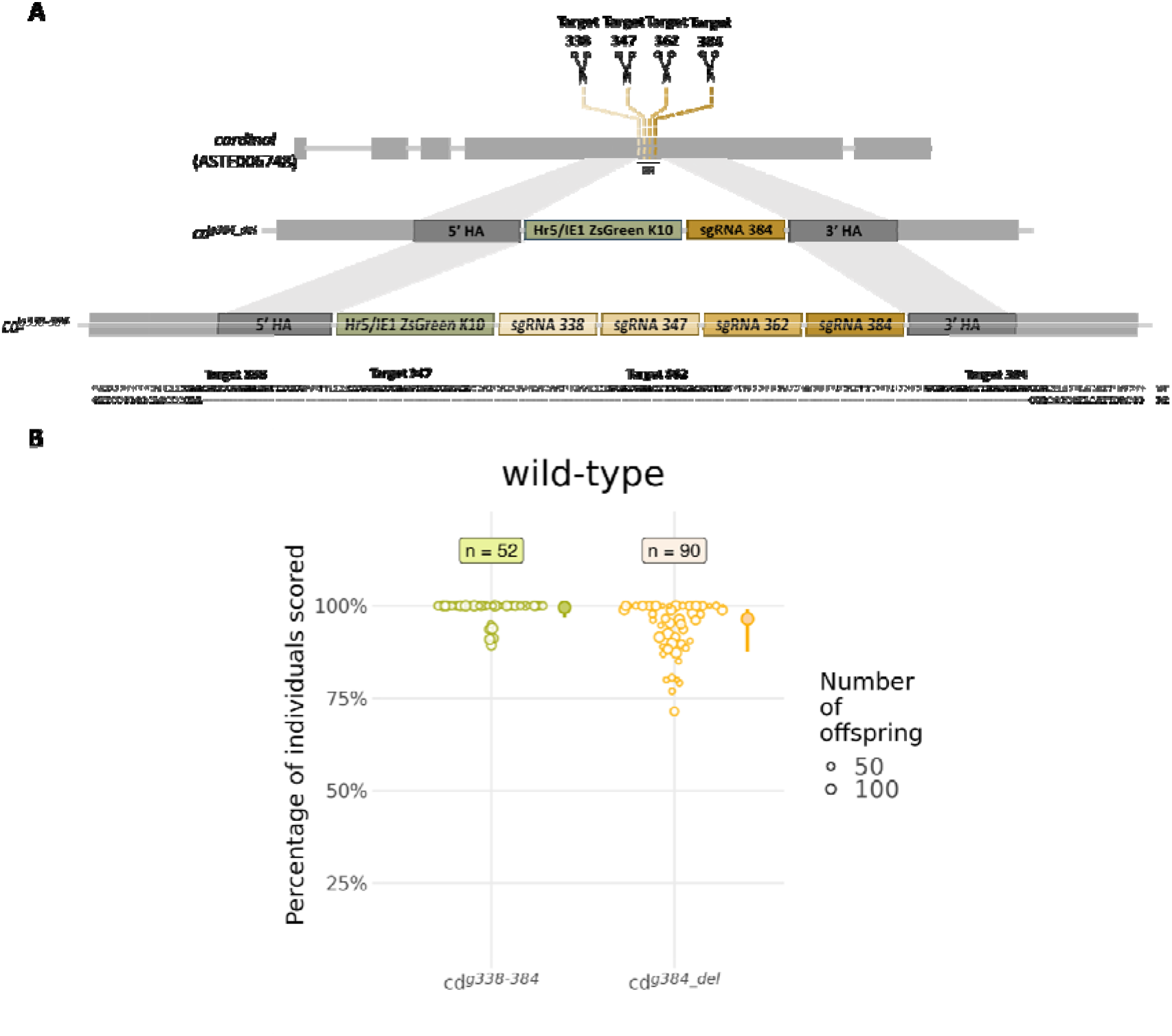
Gene drives using the classical multiplexing strategy have high efficiency. (A) Schematic representation of the *cd* transcript in *An. stephensi*, and the *cd*^*g384_del*^ and *cd*^*g338-384*^ HDR knock-in constructs with their corresponding insertion sites. (B) Inheritance rates of the *cd*^*g384_del*^ and *cd*^*g338-384*^ transgenic lines when crossed to *zpg*^*3’Cas9*^. Circles represent the percentage of F_2_ progeny which inherited the *cd*^*g338-384*^ (green) or the *cd*^*g384_del*^ (yellow) transgenes and the size of the circle is proportional to the number of progeny from that female. Filled circles and error bars represent the mean and the 95% CI, respectively. RR: resected region; n = number of female individuals whose progeny were scored.

To determine drive efficiency, we separately crossed *cd*^*g384_del*^ or *cd*^*g338-384*^ heterozygous males to heterozygous *zpg*^*3’Cas9*^ females to obtain trans-heterozygotes expressing both sgRNA and Cas9. Trans-heterozygous males were then crossed to females which were homozygous mutants for cd (*cd*^*384*^) and their offspring were screened for the presence of the fluorescent markers as well as for eye phenotype. The inheritance rate of *cd*^*g384_del*^ (96.48%; [95% CI] = [87.49-99.08]) (Figure 1B, Table S1) was significantly lower (OR=0.16; z = −2.13; p = 0.033, Table S2) than the inheritance of the same sgRNA but expressed within perfect homology arms, *cd*^*g384*^ (99.43% [98.41-99.8])^34^. This indicates that the requirement for resection during repair does reduce the inheritance rate at this target, here by about 3%. However, this slight reduction in the inheritance rate was reversed by including the 3 additional sgRNAs. Using the same series of crosses we found the *cd*^*g338-384*^ transgene was inherited by 99.55% ([96.87-99.94]) of the progeny (Figure 1B, Table S1), which was not significantly different from the inheritance bias observed for the *cd*^*g384*^ line (OR = 1.26; z = 0.21; p = 0.836, Table S2). No statistically significant differences were observed between the inheritance bias of *cd*^*g384_del*^ and *cd*^*g338-384*^ (OR = 0.12, z = 1.72, p = 0.09, Table S2). We separately reviewed cutting rates, as the frequency of observed mosaicism in the eye colour phenotype, this was observed at 100% for the *cd*^*g338-384*^ transgene (and estimated at >99.9% [99.8-99.9] with a robust glm model), with modest decreases in cleavage rates for the *cd*^*g384*^ line (99.8% [99.7-99.9]; z = −1.99; p = 0.04) and the *cd*^*g384_del*^ line (98.3% [97.8-98.8]; z = −3.48; p < 0.001).

### Inheritance bias is caused by homing and not meiotic drive

To verify that the inheritance bias observed for the *cd*^*g384*^ (data published in ^34^) and the *cd*^*g338-384*^ lines was due to homing and not meiotic drive, we crossed *zpg*^*3’Cas9*^ heterozygous females to AGG1928 heterozygous males (F_0_) to obtain *zpg*^*3’Cas9*^;AGG1928 trans-heterozygotes (F_1_). The AGG1928 transgene, which was generated by piggyBac-mediated random integration, was found to be linked to the cd gene and contains a DmAct5C-ZsYellow-P10 marker cassette as well as other components not relevant for this study. In this experiment, *zpg*^*3’Cas9*^;AGG1928 trans-heterozygous males were crossed to *cd*^*g384*^ or *cd*^*g338-384*^ females, and the remaining crosses performed the same as described above. If meiotic drive occurs we would expect to see a decrease in the inheritance of this linked marked allele from Mendelian inheritance (50%). Despite an inheritance rate of 93.3% ([95% CI] = [92.1-94.2]) for *cd*^*g384*^ and 97.4%([95% CI] = [96.6-98.1]) for *cd*^*g338-384*^, we found no evidence for meiotic drive in the inheritance rate of AGG1928 in either *cd*^*g338-384*^ (49.6%; [95%CI] = [47.2-52]) or *cd*^*g384*^ (48.3%; [46.2-50.5]) (GLMM LRT: χ^2^_1_ = 0.61, p = 0.434; Table S3) crosses.

### Classical multiplexing gives high homing in the presence of a resistance allele

To assess the ability of the classic multiplexing strategy to surpass resistance we generated a Cas9 expressing line which was also homozygous for a 3bp deletion in the sgRNA target site in *cd* (*zpg*^*3’Cas9*^;*cd*^*384*^). Trans-heterozygous *zpg*^*3’Cas9*^;*cd*^*384*^ females were crossed to *cd*^*g338-384*^ heterozygous males (F_0_) to obtain *zpg*^*3’Cas9*^;*cd*^*g338-384/384*^ trans-heterozygous males (Figure 2A), which were then crossed to *cd*^*384*^ females (F_1_). The F_2_ progeny were screened for fluorescence and eye phenotype. We found 96.07% ([95% CI] = [78.43-99.39]) of the F_2_ progeny inherited the *cd*^*g338-384*^ allele (Figure 2B, Table S1). Even though the observed inheritance bias was slightly lower than the inheritance observed in the absence of resistance (99.55% ([95% CI] = [96.87-99.94]), this was not statistically significant (OR = 0.11, z = 1.58, p =0.11, Table S2), showing that the classic multiplexing strategy can sustain high drive efficiency in the presence of a cut-resistant allele. A cross was also performed following the same scheme but using the singleplex *cd*^*g384*^ line (Figure 2A). As expected, the *cd*^*g384*^ transgene was inherited at a Mendelian rate (51.38% [95% CI] = [13.81-87.45]) due to the deletion within the target site of the only sgRNA (Figure 2B).

**Figure 2.**
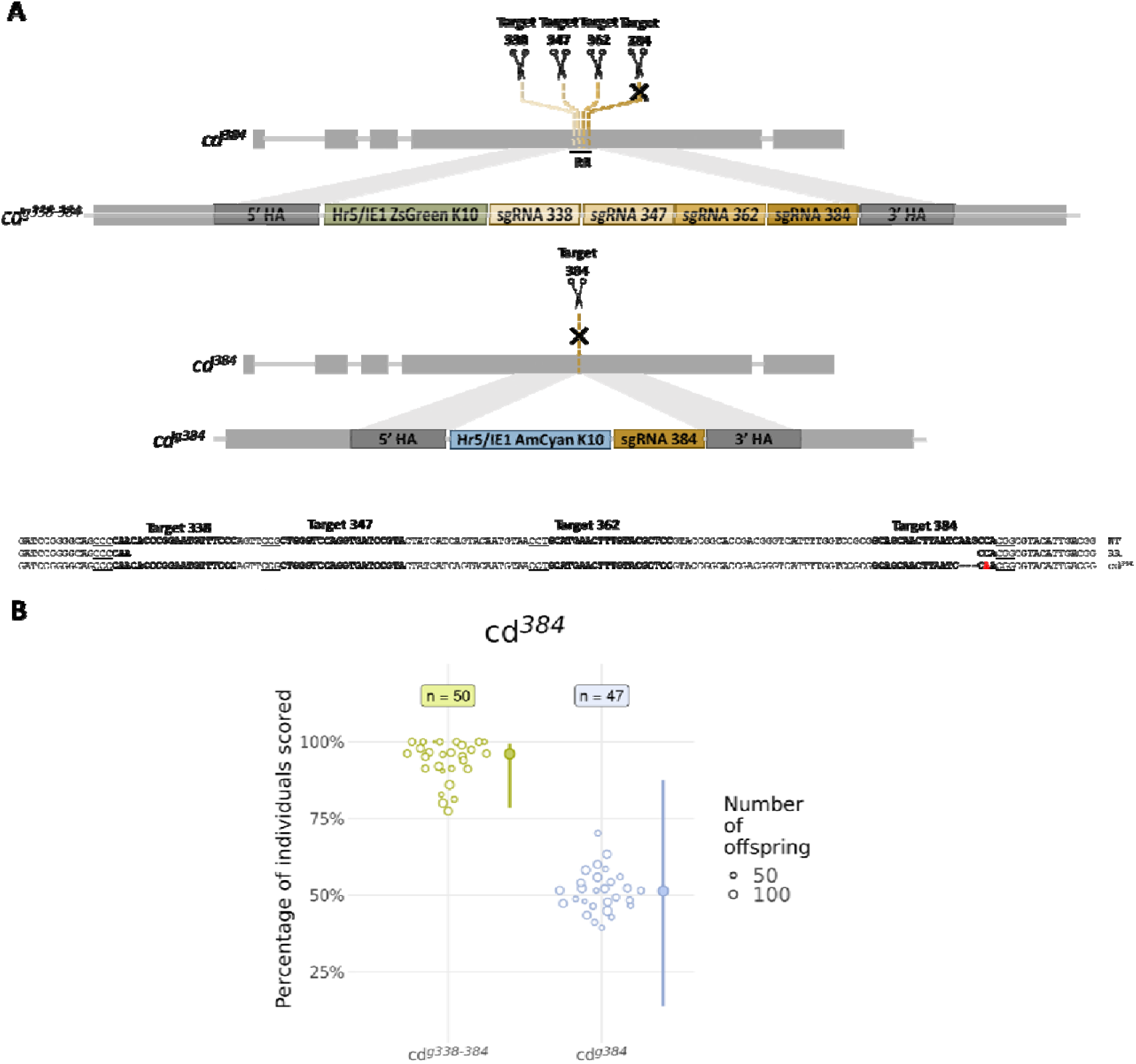
Classical multiplexing can sustain high inheritance rates even in the presence of resistance. Schematic representation of the *cd* transcript in *An. stephensi* including the *cd*^*g338-384*^ and *cd*^*g384*^ HDR knock-in constructs with their corresponding insertion sites (A). The black cross represents the location of the mutation at the 384 cut site. Sequence alignment to illustrate the resected region (RR), and the sequence of the cd384 mutation. (B) Circles represent the percentage of F2 progeny which inherited *cd*^*g384*^ (blue) and *cd*^*g338-384*^ (green) when crossed to *zpg*^*3‘Cas9*^ in the presence of a mutation at the 384 cut site. The size of the circle is proportional to the number of progeny obtained from the female. Filled circles and error bars represent the mean and the 95% CI, respectively. n = number of female individuals whose progeny were scored.

### The additive multiplexing strategy maintains homing in the presence of a resistance allele

To validate the additive multiplexing strategy, we generated a transgenic line composed of an sgRNA expressing cassette targeting and inserted into a site upstream of the 384 target site. For this strategy to be successful the distance between the target sites must be sufficient such that both transgenes can home independently while reducing the probability of segregation via recombination. Therefore, we selected a second target 475 nucleotides away from the 384 target, *cd*^*g225*^ (Figure 3A). The *cd*^*g225*^ line was then crossed to *zpg*^*3’Cas9*^ (F_0_) to determine the homing and cutting rates at the 225 target site. We found 92.08% ([95% CI] = [63.12-98.75]) of the F_2_ progeny inherited the *cd*^*g225*^ allele, indicating that the inheritance rate (OR = 0.07; z = −2.44; p = 0.015) and cleavage rate (95.8% [95.01-96.43]; z = −14.57; p < 0.001, Table S2) at the 225 cut site, while high, are significantly lower than the rate obtained at the 384 site (Figure 3C, Table S1).

**Figure 3:**
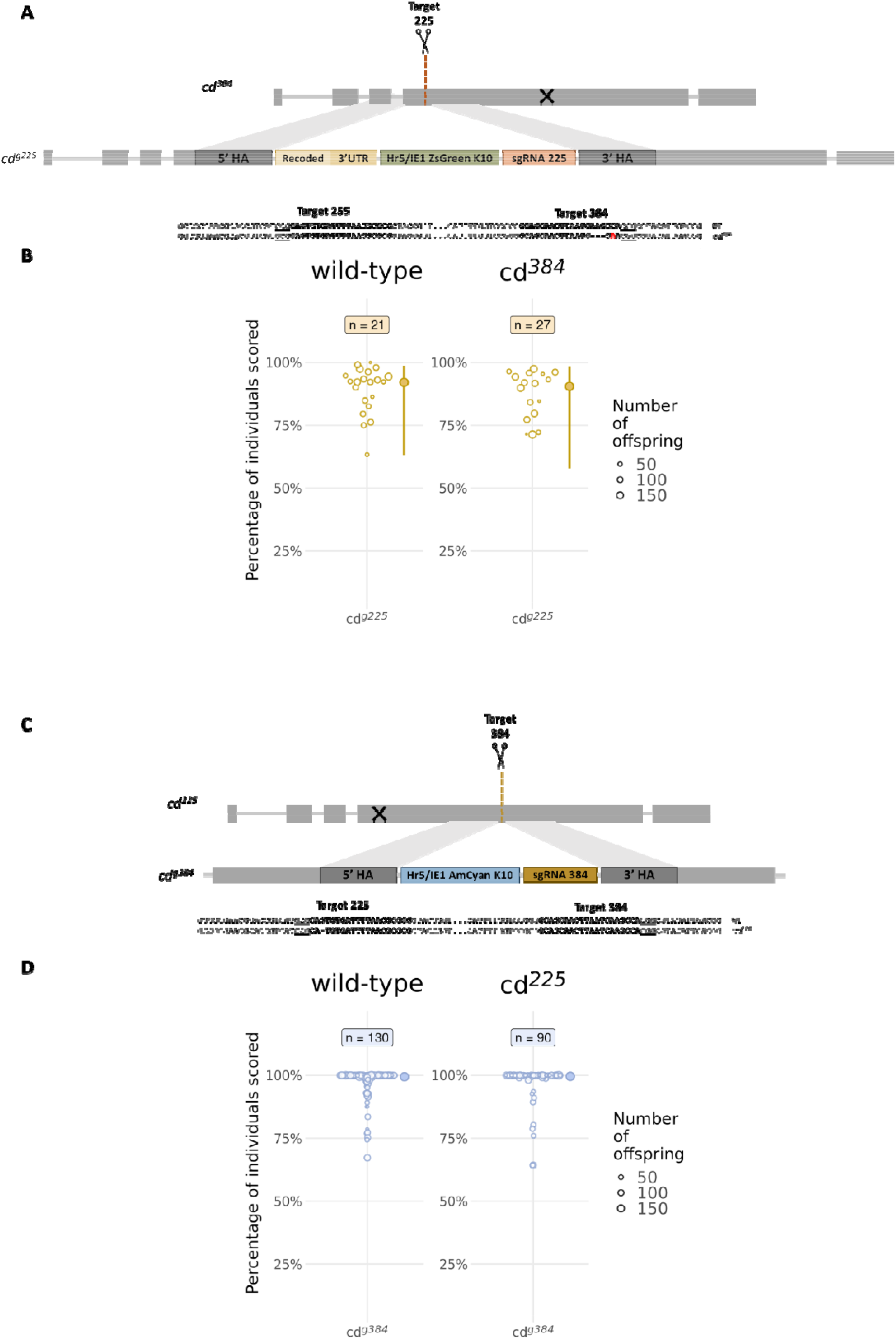
Additive multiplexing maintains high inheritance rates in the presence of pre-existing mutant alleles. Schematic representation of the *cd*^*g225*^ (A) and the *cd*^*g384*^ (C) transgenes and their insertion sites located within the cd transcript and sequence alignment of the cd mutants. The dashed line with the scissors represents the cut site and the black x indicates the location of the resistant mutation in the background of *zpg*^*3‘Cas9*^. (B) Inheritance rates of *cd*^*g225*^ in a wild-type background (left) or in the background of a resistance allele (right). (D) Inheritance rates for the *cd*^*g384*^ allele in a background with (right) or without (left) a deletion affecting the contiguous target site. Circles represent the percentage of F2 offspring that inherited the *cd*^*g225*^ (B) or the *cd*^*g384*^ transgene (D). Data for *cd*^*g384*^ in a wild-type background presented here was originally published in 34. Filled circles and error bars represent the mean and 95% CI. n = number of female individuals whose progeny were scored. The size of the circle is proportional to the number of progeny obtained from the female.

We also wanted to assess the homing and cutting rates of cd^*g225*^ or *cd*^*g384*^ in the presence of a resistant allele at the distant target site (Figure 3A and Figure 3B, Table S1) to determine the behaviour of a drive in a population with resistance alleles to a previously released gene drive. For this purpose, we generated an additional strain of the *zpg*^*3’Cas9*^ line to be homozygous for a resistance allele at the 225 sgRNA target site (*zpg*^*3’Cas9*^;*cd*^*225*^). Female *zpg*^*3’Cas9*^;*cd*^*384*^ were crossed to *cd*^*g225*^ males and *zpg*^*3’Cas9*^;*cd*^*225/225*^ females were crossed to *cd*^*g384*^ males (F_0_) to obtain trans-heterozygotes. Then, trans-heterozygous males for the *zpg*^*3’Cas9*^;*cd*^*384*^ and *cd*^*g225*^ alleles and for the *zpg*^*3’Cas9*^;*cd*^*225*^ and *cd*^*g384*^ alleles were separately crossed to *cd*^*384*^ and *cd*^*225*^ females (F_1_), respectively. F_2_ offspring were screened for fluorescence expression as well as eye phenotype. The *cd*^*g225*^ transgene was inherited by 90.5% ([95%] = [57.88-98.51]) of the F_2_ offspring (Figure 3C, Table S1), which was slightly lower but not significantly different from the inheritance rates in the absence of resistance (OR = 1.22; z = 0.14; p = 0.89, Table S2). Similarly, the *cd*^*g384*^ allele was inherited by 99.59% ([95% CI] = [98.31-99.9]) of the F_2_ progeny (Figure 3D, Table S1). These results suggest that multiple releases of independent single target CRISPR/Cas9-based gene drives could spread in a population in the presence of resistance.

### Deep sequencing reveals variable sgRNA efficiency and generation of resistance alleles

To determine the spectrum of potential resistance alleles generated by the *cd*^*g338-384*^ multiplexed line, we collected F_1_ *zpg*^*3’Cas9*^;*cd*^*g338-384*^, *zpg*^*3’Cas9*^;*cd*^*g384*^ and *zpg*^*3’Cas9*^;*cd*^*g225*^ trans-heterozygous males for Illumina-based amplicon sequencing of the cd target sequence and analysis using CRISPResso2. The sgRNA384 expressed by *zpg*^*3’Cas9*^;*cd*^*g384*^ trans-heterozygous males had the highest mutation rate of all the sgRNAs analysed (63%, Figure 4B). This sgRNA also showed the highest mutation rate (46.1%) in the *zpg*^*3’Cas9*^;*cd*^*g338-384*^ F_1_ trans-heterozygous males while the other three sgRNAs showed relatively lower mutation rates (22.5-26%) at their respective target sites (Figure 4A). sgRNA225 had one of the lowest overall indel rates (26%, Figure 4C). Analysis of wild-type males showed that all nucleotides within the analysis window were 95.5% identical to the reference sequence, with the remaining reads mostly modified by single nucleotide substitutions (SNPs, 97.7% of the modified reads, Figure S1) which we exclude from our analysis. Manual analysis of sequences with indels revealed approximately 75% of indels were out-of-frame in all the transgenic lines (Table S4), indicating that although drive resistant alleles are generated, they are likely to be non-functional. The *zpg*^*3’Cas9*^;*cd*^*g384*^ had the highest proportion of out-of-frame indels (78.5%), compared with that observed in the *zpg*^*3’Cas9*^;*cd*^*g338-384*^ [70.1%] and *zpg*^*3’Cas9*^;*cd*^*g225*^ (71.7%).

**Figure 4:**
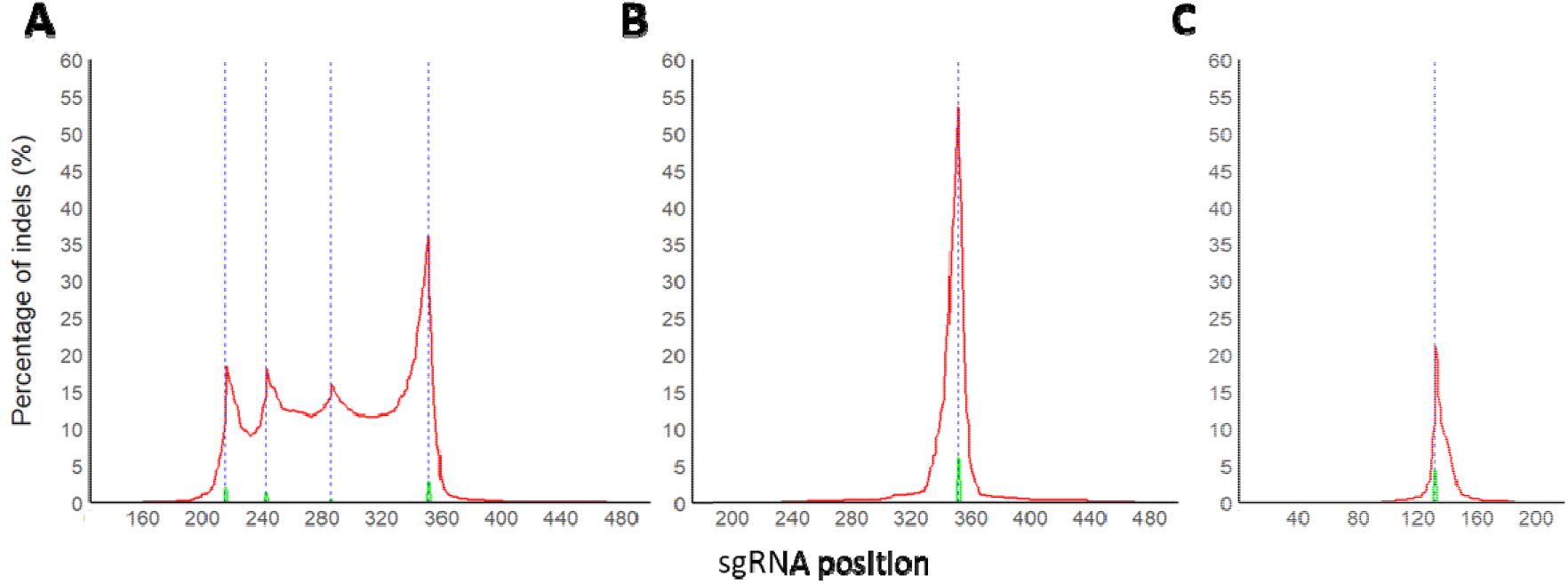
Rates of insertions and deletions (indels) in the target sites of *cd*^*g338-384*^, *cd*^*g384*^and *cd*^*g225*^. Blue dashed lines represent the cut sites of the different sgRNAs. Red line represents the rate of deletions while green line represents the rate of insertions in the *zpg*^*3‘Cas9*^;*cd*^*g338-384*^ (A), the *zpg*^*3‘Cas9*^;*cd*^*g384*^ (B), and the *zpg*^*3‘Cas9*^;*cd*^*g225*^ (C) F1 trans-heterozygous males determined by CRISPResso2. Source data is provided as a Source Data file.

Further analysis of the reads in *zpg*^*3’Cas9*^;*cd*^*g338-384*^ trans-heterozygotes reveals 18% of total reads (30% of modified reads) lack just the sgRNA384 target site and 8.5% of total reads (14% of modified reads) lack all four sgRNA target sites (Figure 5). Of the reads lacking all four target sites, 68% had a single deletion which ablated the target sites. This suggests that simultaneous cutting of the two outermost sgRNAs, 338 and 384, can directly eliminate all four sgRNA targets and create a fully cut-resistant allele in a single step.

**Figure 5.**
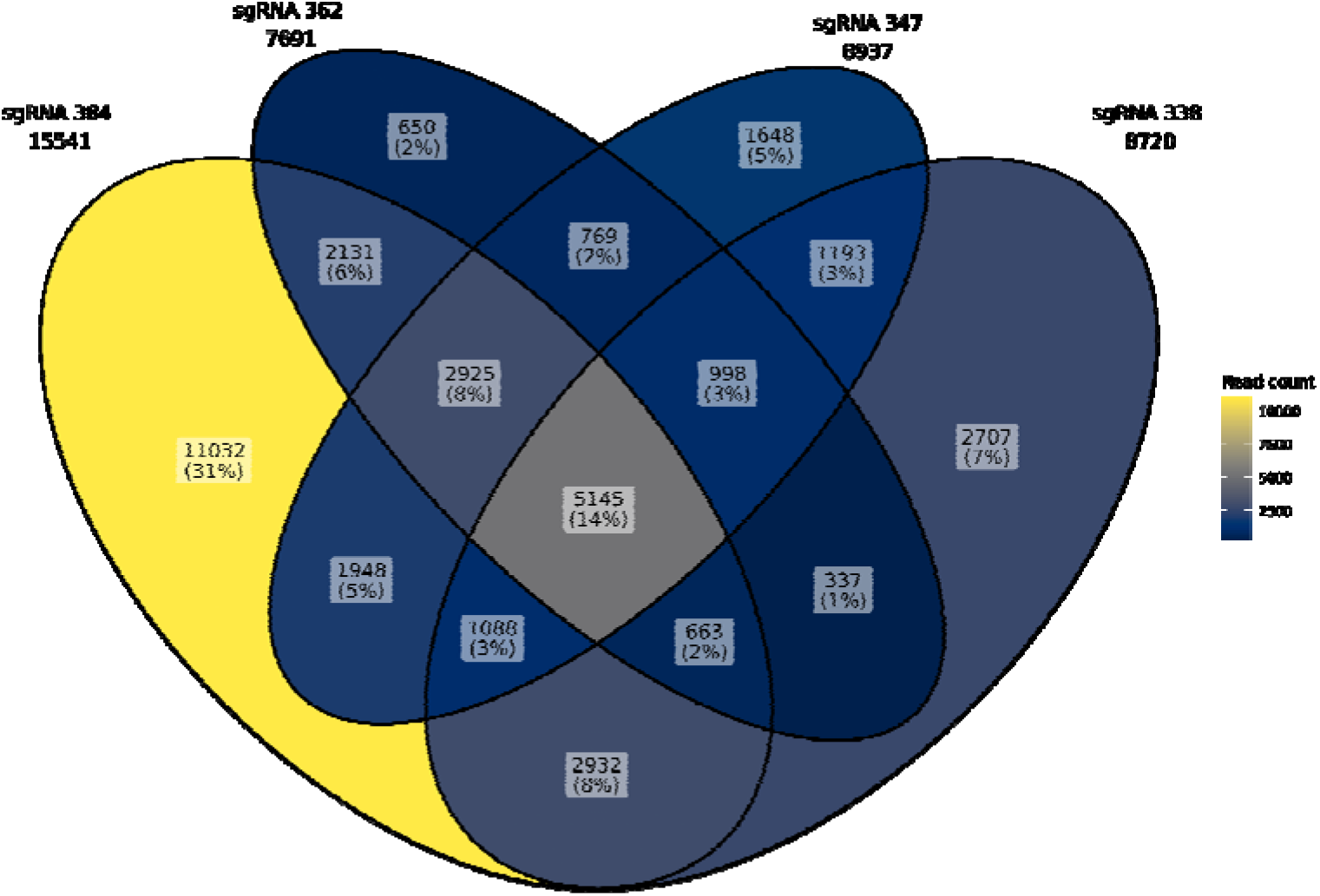
Venn diagram showing the proportion and number of modified reads lacking each sgRNA including all possible combinations. Source data is provided in Source Data File.

Of the ten most common mutations, two have a large deletion that removes all four sgRNA targets, another two have deletions which remove three targets and the remaining six have mutations only at the 384 target site (Figure S2).

### Modelling drive behaviour

The above work demonstrated the potential of genetic elements for both classical and additive multiplexing approaches to achieve high levels of sgRNA target site cleavage and gene drive inheritance. Thus, we use parameters derived here to inform mathematical models, allowing us to further explore the potential ability of these approaches to overcome issues of gene drive resistance and fitness costs alongside their ability to provide lasting population modification.

Mathematical models used here are analogous to those previously used to compare a range of multiplexing approaches, discussed in depth previously^23^. Briefly, the models used here are individual-based and aim to capture a cage-trial type experiment; i.e. relatively low numbers of individuals, non-overlapping generations and no outside ecological factors such as density dependence. Within these models stochastic effects are captured through either random selection (e.g. choosing individuals to seed the next generation and allocation of offspring genotypes) or comparing values randomly chosen from a distribution to selected parameter values (e.g. fitness costs, target site cleavage and repair via homing or NHEJ). We also consider a number of modelling assumptions that are vital in moving from the values derived here to models of full classical or additive multiplexing approaches. Prior experience shows that the majority of gene drive elements impart a fitness cost on carrying individuals, even when inserted into putatively neutral sites. Thus, here we assume each gene drive element has a moderate fitness cost (ε) of 5% relative to wild type, applied multiplicatively such that heterozygote fitness is (1-ε) and homozygote fitness (1-ε) ^2^. Over time the additive approach allows for the accumulation of multiple drive elements in a single allele. Thus as a base assumption we cap the maximum fitness cost imparted on an individual at (1-ε) ^2^ since the elements should be located close enough as to only disrupt a single genomic region. Our other main assumption centres on the rates of sgRNA target cleavage and drive inheritance for additive drive elements. Each of these elements would in reality have their own rates of target cleavage and inheritance, however for simplicity we assume here that each element (four elements to match the number in the classical approach) has the same drive parameters - considering two different scenarios based on the elements analysed above (*cd*^*g225*^ and *cd*^*g384*^). Full parameter sets are detailed in Materials and Methods.

Due to the stochastic nature of the models used here, we run 100 numerical simulations of the classical multiplexing model and the two additive model parameterisations (for *cd*^*g225*^ and *cd*^*g384*^) for each scenario considered (Figure 6). Importantly, both multiplexing approaches are capable of attaining a high carrier frequency, with all simulations reaching a carrier frequency above 0.9 (i.e. >90% of individuals carry at least one drive copy) within ~10 generations. This holds for both parameterisations of the additive multiplex strategy (*cd*^*g225*^ and *cd*^*g384*^).

**Figure 6:**
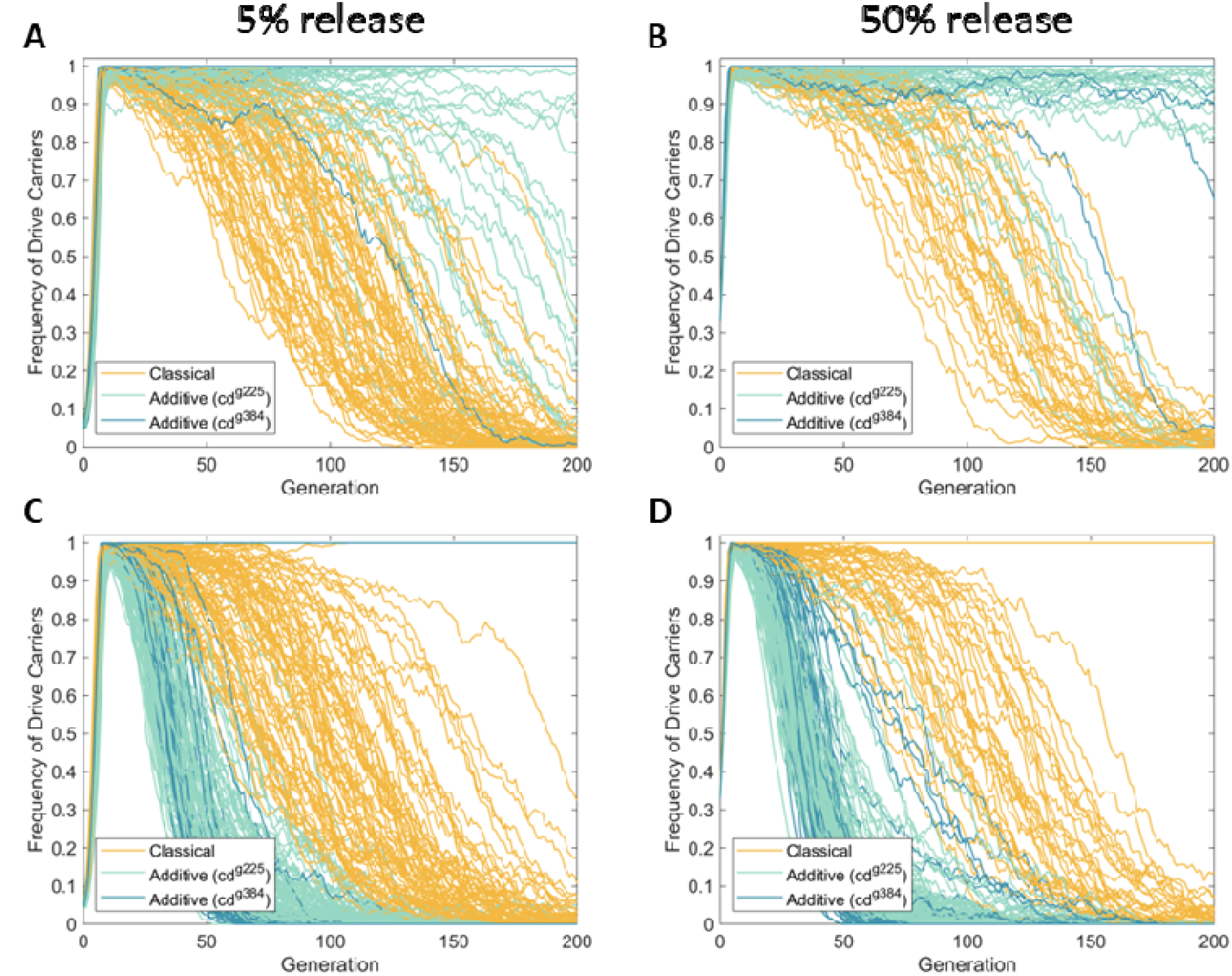
Additive multiplexing can outperform classical multiplexing so long as fitness costs are limited. Sample numerical simulations of the classical multiplexing (yellow) and two parameterisations of the additive multiplexing approach (*cd*^*g225*^ (green) and *cd*^*g384*^ (blue)). The top row (A and B) shows simulations for the scenario in which multiplicative fitness costs for the additive strategy are capped at the value for two copies while the bottom row (C and D) show simulations in which these multiplicative fitness costs are uncapped. The left column (A and C) displays simulations that consider an initial release of 0.05 times the initial wild population whereas the right column (B and D) shows simulations for an initial release of 0.5 times the initial wild population. Note that each line represents a single stochastic numerical simulation and that many lines overlap one another at a drive carrier frequency of one as these represent all of the simulations in which resistance did not take hold.

We first consider the base case in which the multiplicative fitness costs of the additive approach are capped at the value for two drive copies (Figure 6A and B). Here we see that despite the classical and additive approaches reaching high frequency within a similar number of generations, both parameterisations of the additive approach (*cd*^*g225*^ and *cd*^*g384*^) outperform classical multiplexing in terms of their ability to persist within the population. Specifically, for the additive strategy based on *cd*^*g225*^ parameters a total of 10 out of 100 simulations had consistently fallen below a carrier frequency of 0.9 within the 200 generation window of simulation in the large release case, while 19 out of 100 had fallen below this threshold in the small release case. For the additive approach based on *cd*^*g384*^ parameters we saw only 2 and 1 simulations fall below this threshold for the large and small release cases, respectively. This is in stark contrast to the classical multiplexing approach which showed 34 and 76 simulations falling below this threshold for the large and small release cases, respectively with the vast majority of those dropping to near extinction within this time frame. The likely cause of this, as discussed previously^23^ is that, when all (or at least the outermost) sgRNAs cut simultaneously, the system has the potential to repair the region via NHEJ, resulting in the deletion of all target sites within a single generation. Within our previous study^23^, this was found to be the one of the most common causes of drive failure within the classical multiplexing approach. The additive approach, however, does not allow for such an event to occur, likely meaning that accumulation of resistance alleles is far slower, resulting in the improved performance (Figure 6A and B).

Thus far we have considered a base scenario in which the fitness costs of the additive multiplexing approach is capped at that for two drive copies. This appears a reasonable assumption since each additive drive element should be located close enough together that they only disrupt one genomic region. However, another possibility is that by increasing the amount of disruption to that target region through the insertion of multiple additive drive elements may cause fitness costs to increase. Thus, we now progress to consider a scenario in which each additive drive element contributes to the multiplicative fitness cost (i.e. we remove the two element cap on fitness costs). Results of this represent a worst case scenario in terms of the fitness costs imposed by an additive multiplex approach (Figure 6C and D). These results clearly show that the additive approach can be substantially limited if multiple additive drive elements contribute equally to the overall fitness cost. For instance, removing the two element cap on fitness costs causes the number of simulations falling consistently below a drive carrier frequency of 0.9 within 200 generations to rise from 2 to 35 and 1 to 33 for the *cd*^*g384*^ parameterisation in the large and small release scenarios, respectively. Similarly, for the *cd*^*g225*^ parameterisation, the number of simulations falling below this threshold rises from 10 to 100 (i.e. all simulations) and 19 to 100 (i.e. all simulations) for the large and small release cases, respectively. Perhaps more importantly, removing the cap on fitness costs for the additive approach causes it to perform significantly worse than the classical multiplexing approach.

Put together, these results indicate that both the classical and additive multiplexing approaches have the potential to achieve rapid increases in frequency and a good degree of persistence within the population. While the additive approach can overcome a major issue of the classical approach (deletion of all target sites in a single generation), if each additive drive element contributes significantly to the overall fitness cost of carrier individuals then the performance of this approach drops off drastically. Thus, these approaches show significant potential for further development, though further work is needed to fully understand the fitness characteristics of the additive approach.

## Discussion

Strategies that aim to overcome resistance are critical for the success of homing-based gene drives as vector control strategies. Multiplexing has emerged as a promising way to mitigate target site resistance by targeting multiple loci simultaneously^23,24,35–38^. In this study, we compare two multiplexing strategies to a single sgRNA drive and determine their homing and cleavage efficiencies in An. stephensi, being the first time these strategies have been investigated in depth in this mosquito species.

Using an endogenous zpg-driven Cas9, the engineered classical multiplexing line (*cd*^*g338-384*^) showed over 99% germline cutting and inheritance bias, which is slightly, but not significantly, higher than the cutting and homing rates obtained with the singleplex drive (*cd*^*g384*^). Even though the *cd*^*g384_del*^ transgene utilises the most active sgRNA, this line showed the lowest cleavage rate and inheritance bias associated with an increased variance, potentially due to the requirement for resection for all homing events in this line (can be seen in the confidence intervals in Figure 1B). These results suggest the addition of multiple sgRNAs are responsible for the increase in inheritance despite the required resection of the homology arms, which appears to increase the variability of cutting within individuals in the absence of additional sgRNAs. One of the potential challenges of the classical multiplexing strategy is when simultaneous double standard breaks at multiple sites are repaired via NHEJ, resulting in the deletion of the intervening region and creating resistance for all the target sites found within such deletion and potentially to the drive. Analysis of indel formation in trans-heterozygotes revealed that 14% of mutated sequences had lost all four sgRNA target sites under our classical multiplexing design. These rates are higher than previously reported in *Ae. aegypti*^39^, where no complete sgRNA deletions were observed in individuals carrying multiplexed sgRNA expression constructs, so our results represent a worst-case scenario, providing an upper-bound estimate of potential indel formation in the progeny. It should be noted that this experiment sequenced indels as a result of somatic cutting and not germline, but we expect the same range of mutations to be generated. These data all highlight that resistance remains a critical consideration, and alternatives to the classical multiplexed gene drive systems are needed.

To overcome this limitation, alternative multiplexing strategies, including the additive multiplexing strategy, have been proposed^23^. Here, we assessed the efficiency of the additive multiplexing strategy to overcome resistance. Our results highlight the effectiveness of an additive multiplexing strategy, in which each singleplex drive element retains high inheritance rates, regardless of the presence of resistance alleles at the alternate target site(s). This redundancy offers a significant advantage over classical multiplexing strategies that could enable simultaneous cleavage at multiple sites within a single construct, where loss of one or more target sites can compromise drive efficiency. By distributing drive activity across separate loci, the additive design reduces the selective pressure for any one resistant allele to block transmission entirely. Importantly, the consistent performance of each singleplex element both in the presence and absence of resistance suggests that this strategy can provide enhanced stability and long-term efficacy, allowing for iterative releases to overcome any resistance alleles in a given population. Another alternative strategy proposed is the ‘blocking’ strategy^23^, similar in design to the *cd*^*g384_del*^ transgenics generated as a control for this study. In this strategy each construct expresses a single sgRNA and removes (blocks) adjacent target sites. Use of this strategy would eliminate the potential downfall of the classic strategy where a simultaneous cut at the outer two sgRNA targets yields an allele which is resistant to all sgRNAs in a single step. The potential downside to this strategy is the deletion of homologous sequences adjacent to the target site such that all cuts require resection for homing to occur. In D. melanogaster this has been shown to decrease drive efficiency, in some cases completely^29^. However, the high homing rates found with the *cd*^*g384_del*^ *l*ine, with only a 3% decrease from a construct expressing the same sgRNA but with perfect homology arms, indicate this may be different between species and that this strategy is feasible in *An. stephensi*.

Mathematical modelling shows that both classical and additive multiplexing strategies demonstrate considerable promise for achieving high drive frequencies and persist within target populations, with the additive design presenting a slight advantage when fitness costs are capped at two alleles. However, our modeling also underscores a key limitation of the additive multiplexing strategy: its performance is highly sensitive to the cumulative fitness costs imposed by each individual drive element. If each allele contributes substantially to fitness reduction, the combined burden can significantly impede the spread and persistence of the drive in the target population. This trade-off reflects a broader challenge in gene drive development, where the balance between drive efficacy and host fitness must be carefully optimised^16,39–41^. In contrast, classical multiplexing systems may impose lower cumulative costs by packaging multiple gRNAs within a single construct, though they remain vulnerable to large resistance-inducing deletions. Therefore, the choice between multiplexing strategies may depend on the specific biological and ecological context, including target species, population structure, and regulatory constraints. Importantly, future work should focus on experimentally quantifying the fitness costs associated with multi-element additive drives, particularly in heterozygous individuals, as well as their stability over multiple generations. Overall, both multiplexing strategies offer promising avenues for enhancing the robustness of homing-based gene drives, and further refinement of these approaches will be crucial to developing safe, effective, and confinable tools for the control of An. stephensi and other mosquito vectors of disease.

## Material and methods

### Maintenance of mosquito colony

An. stephensi of the SDA-500 wild-type strain and transgenics were reared under standard conditions of 70-80% relative humidity, 28±1°C, and 14:10h day-night cycle^42,43^. First instar larvae were fed with Sera Micron (Olibetta) and with Extra Select Pond Pellets Complete Fish Food at later stages. Adults were fed with 10% sucrose ad libitum and females were blood-fed with defibrinated horse blood (TCS Biosciences) administered through a Hemotek membrane feeding system (Hemotek, Inc) covered with a double layer of Parafilm (Bemis).

### Plasmid design and synthesis

AGG2072 (*cd*^*g384_del*^) was synthesised and contains ~1.5kb homology arms, the Hr5/IE1-ZsGreen fluorescent marker and the As7SK regulatory regions^34^ expressing the sgRNA (GCAGCAACTTAATCAAGCCA).

AGG2301 (*cd*^*g338-384*^) was synthesised starting with the AGG2072 plasmid and adding 3 additional sgRNA expression cassettes utilising three previously characterised pol III promoters^34^ to express the additional sgRNAs (U6A-GGAGCGTACAAAGTTCATGC, U6B-TACGGATCACCTGGACCCAG and U6C-GGGAAACATTCCGGGTGTTG). It should be noted that due to their proximity in the genome the region used in the reference for the U6A terminator overlaps with the U6B promoter region, this region has not been duplicated in the construct to avoid the potential for recombination. To avoid repetition of the sgRNA backbone, alternatives were selected based on a pilot experiment in Aag2 cells (Figure S2).

AGG2273 (*cd*^*g225*^) was synthesised and consists of ~1kb homology arms, *An. gambiae cd* (AGAP003502) was used to replace cd coding sequence followed by a recoded ex6 of *An. stephensi cd*, and the AGAP003505 3’UTR. A cassette to express the sgRNA (CGCGCGTTAAAATCACACTG) consists of the An ste U6C regulatory regions^34^. An additional cassette contains the Hr5/IE1 promoter expressing the ZsGreen fluorescent marker protein. Due to poor fitness of this line, a second version of this construct was also made without the recoding (AGG2360).

Complete plasmid sequences are available through NCBI: AGG2072_cdg384_del: PV342346, AGG2360_cdg225:PV342348, AGG2301_cdg338-384:PV342349.

### Microinjection and screening of transgenic mosquitoes

Transgenic mosquitoes were generated by microinjection of embryos following a previously described protocol^42,43^. Briefly, adult mosquitoes reared under standard conditions but under a reversed light cycle, were blood fed with defibrinated horse blood (TCS Biosciences). 3-5 days after blood feeding, females were allowed to lay eggs in an egg collection cup for 45 minutes. The collected embryos were aligned, transferred into a coverslip using a double sided tape (3M), and covered with halocarbon oil 27 (Sigma). Embryo injections were performed using quartz capillaries (Sutter QF1007010) pulled with a P2000 laser pipette puller (Sutter) and a FemtoJet 4x microinjector (Eppendorf). Embryos were injected with 1X injection buffer^44^, 300ng/⍰l of the donor plasmid (except for AGG2072 800ng/⍰l was used), and 300ng/⍰l of AGG1760 plasmid expressing Cas9 or Cas9 protein (PNABio). The injected embryos hatched approximately 24h after injections and the G_0_ larvae were transferred to rearing trays where they were standard reared. G_0_ adults were pooled according to sex in groups of 20 with 20 G_0_ females crossed to 40 WT males or 20 G_0_ males crossed to 100 SDA-500 females. The offspring (G_1_) were screened for the presence of the fluorescent marker as larvae using a Leica MZ165FC microscope (Leica Biosystems). Positive G_1_ adults were individually outcrossed to SDA-500 to generate isolines. Once each isoline was secured, the G_1_ mosquitoes were collected for gDNA extraction, which was used as a template for PCR and sequence confirmation of the insertion site (Figure S3).

### Generation of *cd*^*225*^and *cd*^*384*^ knock-out lines

Two different cd knock-out lines were generated to determine the cutting rate induced by Cas9. The *cd*^*225*^ knock-out line was generated by crossing *zpg*^3’Cas9^ expressing females to *cd*^*g225*^ heterozygous males^34^. All the non-fluorescent offspring were intercrossed and blood fed. Their offspring were screened for a pink eyed phenotype. The mutation of a single pink eyed female was sequenced, showing a 1bp deletion which results in a frame shift and a stop codon after amino acid 251. The generation of the *cd*^*384*^ knock-out line was previously described^34^, where it was referred to as *cd*^*-/-*^.

### Generation of Cas9 lines with *cd*^*225*^and *cd*^*384*^ background

To determine the homing and cutting rates of the different sgRNA-expressing constructs in the presence of pre-existing target site resistance, fifty *zpg*^3’Cas9^ heterozygous males were crossed with fifty homozygous females of the *cd*^*384*^ and *cd*^*225*^ lines, to generate double heterozygous *zpg*^3’Cas9^;*cd*^*384*^ and *zpg*^3’Cas9^;*cd*^*225*^ lines. Subsequently fifty double heterozygotes were then crossed to homozygous *cd*^*384*^ or *cd*^*225*^ lines. From those progeny, fifty pink eyed males containing the *zpg*^3’Cas9^ transgene were selected and crossed to homozygous *cd*^*384*^ or *cd*^*225*^ females, and the line was maintained by repeating this cross every generation.

### Assessing the CRISPR/Cas9 induced homing and cutting rates

Fifty *zpg*^3’Cas9^ heterozygous females were crossed to fifty heterozygous males of the pertinent cd sgRNA expressing line (F_0_) to obtain trans-heterozygotes. Larvae with both transgene markers were sorted using a Biosorter (Union Biometrica). In addition, 200 non-sorted larvae were counted using the Biosorter. Sorted larvae were maintained under standard conditions and reared to adulthood. Non-sorted larvae were screened as L4 to obtain the ratios of the genotypes generated in the F_1_. Adult male trans-heterozygotes (F_1_) for the transgene expressing the sgRNA and the *zpg*^*3’Cas9*^ (or *zpg*^3’Cas9^;*cd*^*384*^ or *zpg*^3’Cas9^;*cd*^*225*^) allele were crossed in a 1:1 ratio to cd mutant females. The females were allowed to mate for a minimum of 5 days, when they were offered a blood meal. After 24h, females were transferred into disposable espresso coffee cups (Somoplast) containing 30mL of RO water where they laid eggs. F_2_ larvae hatched and were maintained in the coffee cups until they were screened for fluorescence and eye phenotype under a Leica MZ165FC fluorescent stereo microscope (Leica Biosystems) as late larvae. The proportion of F_2_ larvae which inherited the transgene expressing the sgRNA(s) was used to determine the inheritance rates and the proportion of F_2_ larvae which presented a pink eyed phenotype was used to determine the cutting rate for *cd*^*g384_del*^ and *cd*^*g338-384*^. For line (*cd*^*g225*^) due to the recoding, heterozygotes will maintain a WT eye phenotype, so for crossing involving this line the cleavage rate was calculated by adding the rate of inheritance of *cd*^*g225*^ to the number of mosaics in the non-*cd*^*g225*^ inheriting progeny.

### Amplicon sequencing and analysis

Genomic DNA of 10 F_1_ trans-heterozygous males of *zpg*^*3’Cas9*^;*cd*^*g338-384*^, *zpg*^*3’Cas9*^;*cd*^*g384*^, or *zpg*^*3’Cas9*^;*cd*^*g225*^ (AGG2360) and 10 SDA-500 males was extracted using the NucleoSpin Tissue kit (MachereyNagel). A 280bp or 471 bp fragment surrounding the sgRNA target sites of the respective transgenic lines was amplified using primers listed in Table S6. The amplicons were visualized by gel electrophoresis then purified using the NucleoSpin Gel and PCR Clean-up Kit (MachereyNagel) and sent for amplicon sequencing using the Illumina-based AMPLICON-EZ sequencing service from Genewiz (Leipzig, Germany). The sequencing reads obtained were analyzed using CRISPResso2 (Table S7). The indel rates surrounding the sgRNA recognition sites were plotted using ggplot2 in R version 4.4.2.

#### Frameshift Mutation Analysis

In-frame and out-of-frame indels in modified sequences were determined by calculating if the number of nucleotides inserted or deleted is divisible by three using a formula in Microsoft Excel (formula: “=MOD((number of deletions - number of insertions), 3) = 0”). The proportion of frameshift indels in all reads with indels were calculated.

#### Resistance Allele Formation Analysis

The rates of resistance allele formation in *cd*^*g338-384*^ were determined by counting the number of read sequences lacking the sequences of sgRNAs (1, 2, 3, and 4), as well as 3m and 4m (single SNP commonly found in wildtype in sgRNA 3 and 4), using the COUNTIF function in Microsoft Excel. The syntax and datasets used for analysis are available in the Source data file.

### Statistical analysis

Analyses of inheritance ratios were carried out using R version 4.3.3^45^. A power analysis was used to predetermine approximate sample size prior to experiments also using R. Using the parameters 2 groups and an Alpha of 0.01, with an estimated standard deviation of 0.3, the progeny of at least 20 individuals would be necessary to determine a 20% difference in inheritance with a power of 0.8. With this in mind crosses were generated to assess at least this number. Estimated means and 95% confidence intervals were calculated by a Wald z-distribution approximation from a Generalized Linear Mixed Model (GLMM), with a binomial (‘logit’ link) error distribution fitted using the glmmTMB package^46^. Where applicable, initial parameters included the sgRNA transgenic line and the genetic background of the *zpg*^*3’Cas9*^;*cd*^*+/-*^ line; individual replicates were included as a random effect. The model with the best fit was then chosen by comparing Akaike Information Criterion (AIC) and summarized with ‘emmeans’^47^, model residuals were checked for violations of assumptions using the ‘DHARMa’ package^48^. No data were excluded from the analyses. Scripts and raw data can be found at (https://github.com/Philip-Leftwich/stephensii_multiplexing).

### Modelling

Mathematical approaches used here are discussed extensively in our previous work^23^ with details and assumptions specific to this work detailed above. Simulations were conducted using MATLAB (version R2025a; The MathWorks Inc., Natick, MA) while plots were created using a combination of MATLAB versions R2025a and R2023a. Model scripts and associated output data are available via the Open Science Framework (https://osf.io/tsquv/). Parameters used within the model scripts are as follows.

All approaches: Num_Targets = 4, Rel_Ratio = 0.50 or 0.05 (high and low release sizes), Num_Mosquitoes = 500, Num_Eggs = 15, Num_Generations = 200.

Classical: Prob_Cutting = 1, Prob_Home = 0.991, Prob_NHEJ = 1-Prob_Home, RelFit_Het = 0.95, RelFit_Hom = 0.9025, RelFit_WT = 1.

Additive (*cd*^*g225*^): Prob_Cutting = 0.958, Prob_Home = 0.8785, Prob_NHEJ = 1-Prob_Home, RelFit_Het = 0.95, RelFit_WT = 1.

Additive (*cd*^*g384*^): Prob_Cutting = 1, Prob_Home = 0.9918, Prob_NHEJ = 1-Prob_Home, RelFit_Het = 0.95, RelFit_WT = 1.

### Data availability statement

All data generated for this manuscript is available in the manuscript, the supplemental files, and the NCBI public database. AmpliconSeq raw data available in NCBI: Bioproject PRJNA1269442. Complete plasmid sequences are available through NCBI: AGG2072_cdg384_del: PV342346, AGG2360_cdg225:PV342348, AGG2301_cdg338-384:PV342349. Chromatograms of Sanger sequencing to confirm the transgene insertions is available at doi: https://doi.org/10.15124/d40a2165-fb3d-4458-9267-b6524858e6a8. Scripts for Crispresso2 analysis, statistical analysis and raw data can be found at (https://github.com/Philip-Leftwich/stephensii_multiplexing). Model scripts and associated output data are available via the Open Science Framework (https://osf.io/tsquv/).

## Acknowledgements

This work was supported, in whole or in part, by funding from the Gates Foundation [INV-008549]. The conclusions and opinions expressed in this work are those of the author(s) alone and shall not be attributed to the Foundation. Under the grant conditions of the Foundation, a Creative Commons Attribution 4.0 License has already been assigned to the Author Accepted Manuscript version that might arise from this submission. Please note works submitted as a preprint have not undergone a peer review process. This work was also supported by strategic funding to The Pirbright Institute from the Biotechnology and Biological Sciences Research Council [BBS/E/I/00007033, BBS/E/I/00007038, and BBS/E/I/00007039].

## Author contributions

MG, EG, LS, SR, JS, KN, JXDA, MAEA performed the crosses and collected the inheritance data. PL performed statistical analysis. MAEA and JTC prepared samples for Amplicon-Seq and EA and JL analysed the data. ME performed mathematical modelling. MAEA, JXDA and LA supervised the work. JMA, AD, MM, EA provided resources and assisted in the insectary. LA procured funding and conceptualised the work.

