## Supplementary information for "Integrating multiplexing into confineable gene drives effectively overrides resistance in *Anopheles stephensi*"

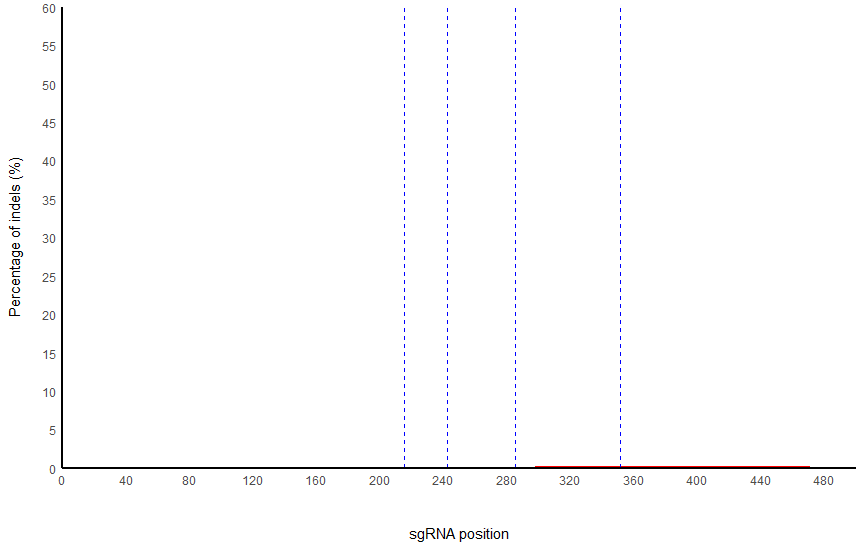

**Figure S1. Rates of insertions and deletions (indels) in the *cd* gene of WT males.** Figure was obtained through CRISPResso2 and data can be found in the Source Data file.

```
GAGTGCTGTGATCCGGGGCAGCCCCAACACCCGGAATGTTTCCCAGTTCCGCTGGGTCCAGGTGATCCGTACTATCATCAGTACAATGTAACCTGCATGAACTTTGTACGCTCCGTACCGGCACCGACGGGTCATTTTGGTCCGCGGCAGCAACTTAATCAAGCCACGGCGTACATTGACGGCTCGGTTGT	Reference
                              sgRNA338                       sgRNA347                                     sgRNA362                                              sgRNA384

GAGTGCTGTGATCCGGGGC-------------------------------------------------------------------------------------------------------------------------------------------------------GTACATTGACGGCTCGGTTGT	0.45% (271 reads)
GAGTGCTGTGATCCGGGGCAGCCCCAACACCCGGAATGTTTCCCCGTTCCGCTGGGTCCGGGTGATCCGTACTATCATCAGTACAATGTAACCTGCATGAACTTTGTACGCTCCGTACCGGCACCGACGGGTCATTTTGGTCCGCGGCAGCAAC----------CACGGCGTACATTGACGGCTCGGTTGT	0.40% (244 reads)
GAGTGCTGTGATCCGGGGCAGCCCCAA----------------------------------------------------------------------------------------------------------------------------------------CCACGGCGTACATTGACGGCTCGGTTGT	0.34% (203 reads)
GAGTGCTGTGATCCGGGGCAGCCCCAACACCCGGAATGTTTCCCCGTTCCG---------------------------------------------------------------------------------------------------------------------GCGTACATTGACGGCTCGGTTGT	0.28% (170 reads)
GAGTGCTGTGATCCGGGGCAGCCCCAACACCCGGAATGTTTCCCCGTTCCGCTGGGTCCGGGTGATCCGTACTATCATCAGTACAATGTAACCTGCATGAACTTTGTACGCTCCGTACCGGCACCGACGGGTCATTTTGGTCCGCGGCAGCAACTTAA-----CCACGGCGTACATTGACGGCTCGGTTGT	0.26% (158 reads)
GAGTGCTGTGATCCGGGGCAGCCCCAACACCCGGAATGTTTCCCCGTTCCGCTGGGTCCGGGTGATCCGTACTATCATCAGTACAATGTAACCTGCATGAACTTTGTACGCTCCGTACCGGCACCGACGGGTCATTTTGGTCCGCGGCAGCAACTTAATC----CACGGCGTACATTGACGGCTCGGTTGT	0.25% (151 reads)
GAGTGCTGTGATCCGGGGCAGCCCCAACACCCGGAATGTTTCCCCGTTCCGCTGGGTCCGGGTGATCCGTACTATCATCAGTACAATGTAACCTGCATGAACTTTGTACGCTCCGTACCGGCACCGACGGGTCATTTTGGTCCGCGGCAGCAACTTAATCA-----CGGCGTACATTGACGGCTCGGTTGT	0.25% (149 reads)
GAGTGCTGTGATCCGGGGCAGCCCCAACACCCGGAATGTTTCCCCGTTCC-------------------------------------------------------------------------------------------------------------------ACGGCGTACATTGACGGCTCGGTTGT	0.25% (148 reads)
GAGTGCTGTGATCCGGGGCAGCCCCAACACCCGGAATGTTTCCCCGTTCCGCTGGGTCCGGGTGATCCGTACTATCATCAGTACAATGTAACCTGCATGAACTTTGTACGCTCCGTACCGGCACCGACGGGTCATTTTGGTCCGCGGCAGCAAC-------------GGCGTACATTGACGGCTCGGTTGT	0.25% (148 reads)
GAGTGCTGTGATCCGGGGCAGCCCCAACACCCGGAATGTTTCCCCGTTCCGCTGGGTCCGGGTGATCCGTACTATCATCAGTACAATGTAACCTGCATGAACTTTGTACGCTCCGTACCGGCACCGACGGGTCATTTTGGTCCGCGGCAGCAACTT-------CCACGGCGTACATTGACGGCTCGGTTGT	0.22% (135 reads)
```

**Figure S2. Ten most frequent mutations in *cd^g338-384^*.** Alignment of the ten most common mutations, with sgRNA target sites underlined and PAM sites highlighted. Proportion of total reads is indicated as a percentage, with the number of reads that had that sequence indicated to the right of each sequence.

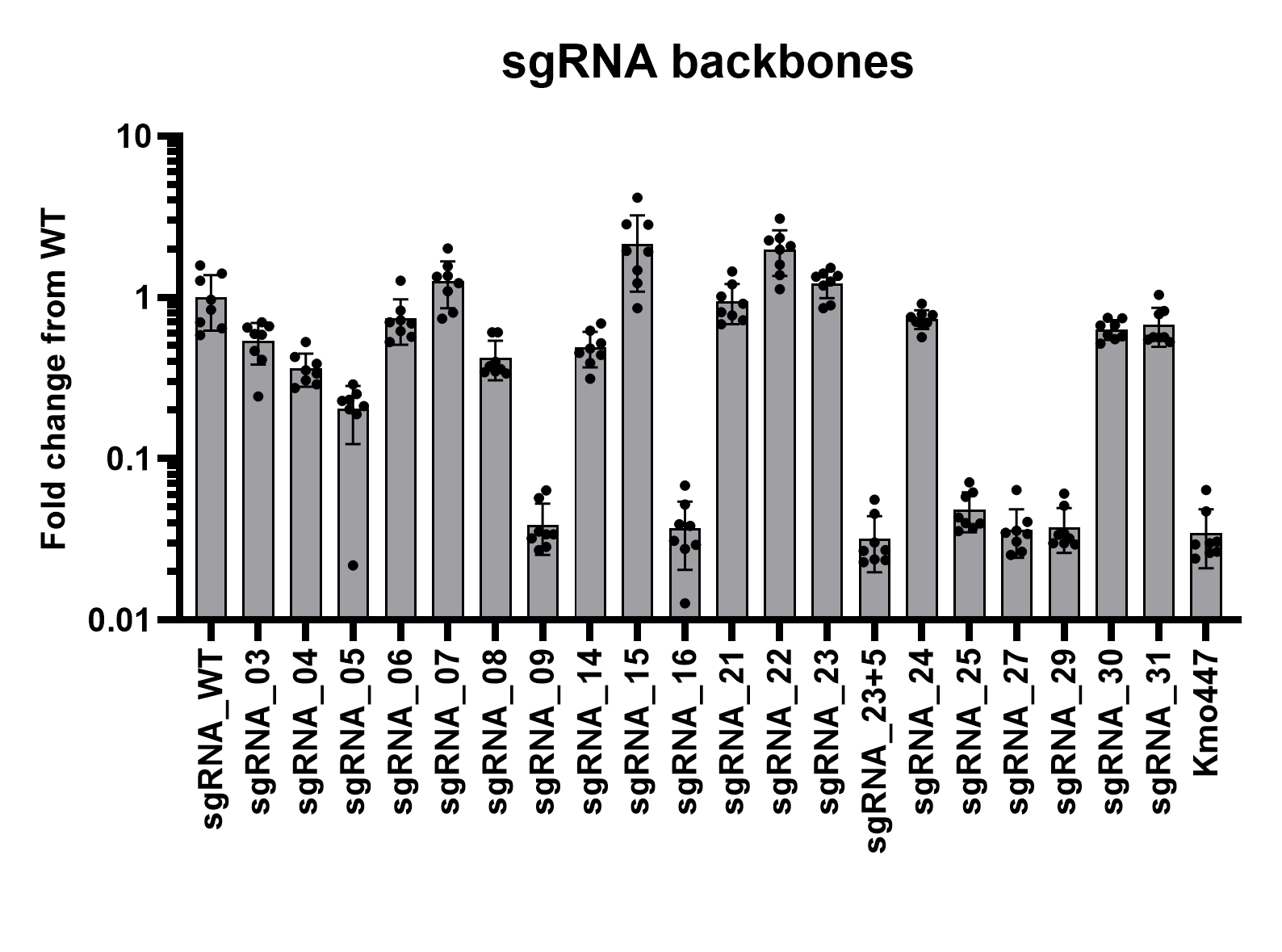

**Figure S3. dCas9-VPR assay** ^1^ **to determine suitability of different sgRNA backbone sequences** ^2^ **in mosquito cells.** Aag2 cells were seeded in a 96 well plate and transfected using TransIT-Pro (Mirius) with 25ng dCas9-VPR, 25ng TRE-Firefly, 50ng pRL-OpIE2, 40ng *in vitro* transcribed sgRNA ^3^ per well. After 48 hours cells were lysed in 1x Passive Lysis Buffer and the Dual luciferase assay performed on a GloMax Multi+ plate reader (Promega). Ratio of firefly to Renilla luciferase activity is normalised to the WT backbone sequence (sgRNA_WT). Kmo447 does not have a target site present in the reporter plasmid and is used to determine background.

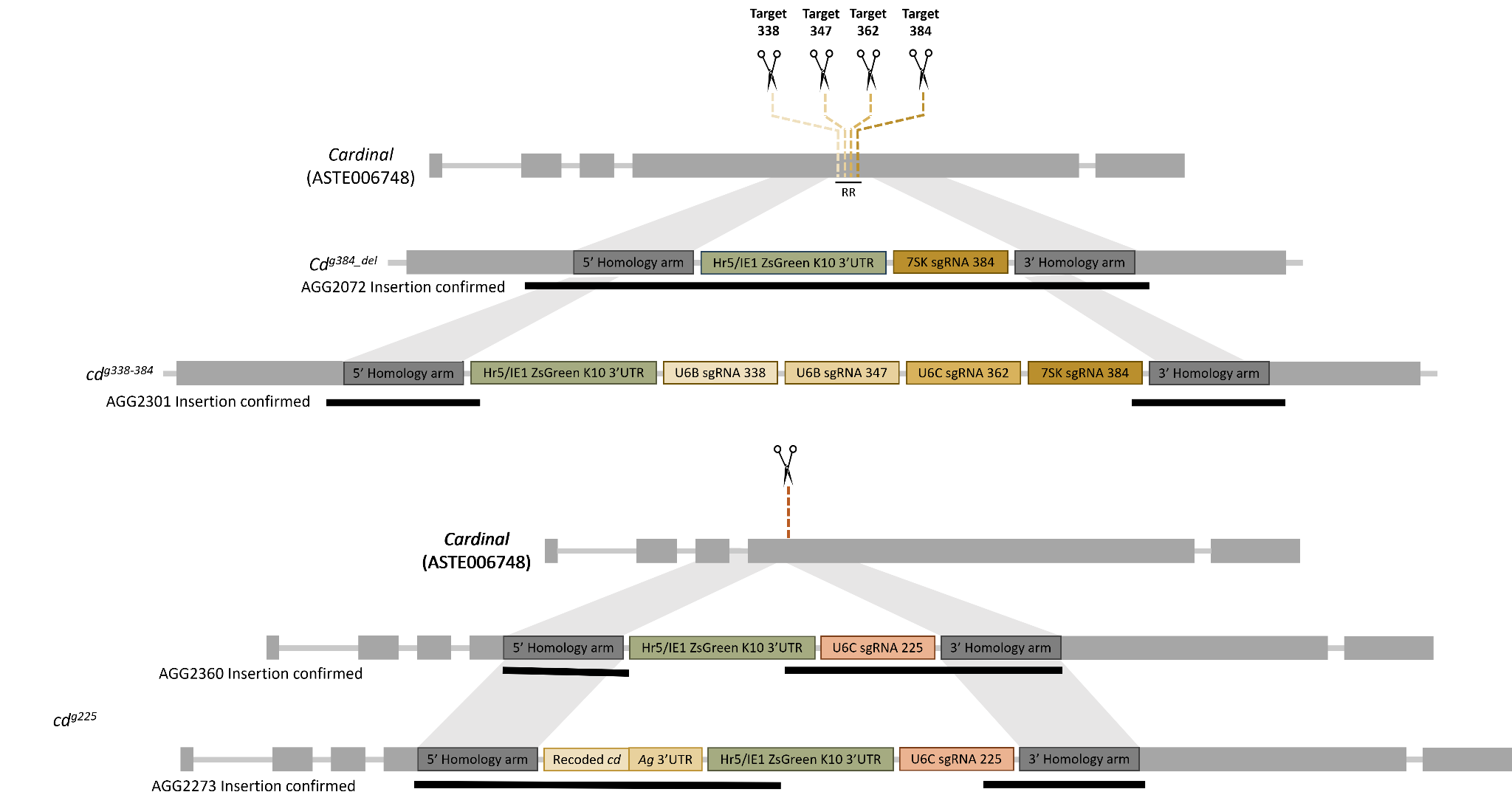

**Figure S4. PCR confirmation of the insertion site for all the generated lines.** Black bars indicate regions which were amplified by PCR and Sanger sequence confirmed. Raw sequences are available at <https://doi.org/10.15124/d40a2165-fb3d-4458-9267-b6524858e6a8>.

**Table S1.** Full model summaries for estimates of inheritance rates from sgRNAs. Log-odds and 95% confidence intervals, and significance values are taken from a binomial glmm with 'logit’ link using sgRNA line and genetic background of the Cas9 bearing parent with full interaction terms. Mixed-effects models included replicate as a random factor.

|  | **Inheritance Rate** | | |
| --- | --- | --- | --- |
| ***Predictors*** | ***Odds Ratios*** | ***95% CI*** | ***P-Value*** |
| (Intercept) | 11.63 | 1.71 – 78.99 | 0.012 |
| gRNA_typecd*g338-384* | 18.95 | 1.22 – 293.69 | 0.035 |
| gRNA type  [cd*g384_del*] | 2.36 | 0.22 – 24.75 | 0.475 |
| gRNA type  [cd*g384*] | 15.02 | 1.71 – 132.34 | 0.015 |
| pre cut  [cd*225*] | 1.40 | 0.24 – 8.01 | 0.708 |
| pre cut  [cd*384*] | 0.82 | 0.05 – 12.49 | 0.886 |
| gRNA_typecd*g338-384*:pre_cutcd*384* | 0.14 | 0.00 – 6.38 | 0.309 |
| gRNA type  [cd*g384*]  × pre cut  [cd*384*] | 0.01 | 0.00 – 0.24 | 0.006 |
| **Random Effects** | | | |
| σ2 | 4.93 | | |
| τ00 id:group_letter | 1.64 | | |
| τ00 group_letter | 0.86 | | |
| ICC | 0.15 | | |
| N id | 328 | | |
| N group_letter | 13 | | |
| Observations | 328 | | |
| Marginal R2 / Conditional R2 | 0.325 / 0.425 | | |

**Table S2.** Pairwise contrasts (with tukey adjusted P-values) for all sgRNA lines and genetic backgrounds derived from a fitted generalized binomial mixed-effects model (see STable 1).

| **contrast** | **odds.ratio** | **SE** |  | **z.ratio** | **p.value** |
| --- | --- | --- | --- | --- | --- |
| (**cd*g225* wild-type) / (cd*g338-384* wild-type)** | 0.05 | 0.07 |  | -2.1 | 0.04 |
| **(cd*g225* wild-type) / (cd*g384_del* wild-type)** | 0.42 | 0.51 |  | -0.71 | 0.48 |
| **(cd*g225* wild-type) / (cd*g384* wild-type)** | 0.07 | 0.07 |  | -2.44 | 0.01 |
| **(cd*g225* wild-type) / (cd*g384* cd*225*)** | 0.05 | 0.06 |  | -2.49 | 0.01 |
| **(cd*g225* wild-type) / (cd*g225* cd*384*)** | 1.22 | 1.7 |  | 0.14 | 0.89 |
| **(cd*g225* wild-type) / (cd*g338-384* cd*384*)** | 0.48 | 0.66 |  | -0.54 | 0.59 |
| **(cd*g225* wild-type) / (cd*g384* cd*384*)** | 11 | 15.09 |  | 1.75 | 0.08 |
| **(cd*g338-384* wild-type) / (cd*g384_del* wild-type)** | 8.04 | 9.77 |  | 1.72 | 0.09 |
| **(cd*g338-384* wild-type) / (cd*g384* wild-type)** | 1.26 | 1.42 |  | 0.21 | 0.84 |
| **(cd*g338-384* wild-type) / (cd*g384* cd*225*)** | 0.9 | 1.11 |  | -0.08 | 0.93 |
| **(cd*g338-384* wild-type) / (cd*g225* cd*384*)** | 23.12 | 32.51 |  | 2.23 | 0.03 |
| **(cd*g338-384* wild-type) / (cd*g338-384* cd*384*)** | 9.03 | 12.56 |  | 1.58 | 0.11 |
| **(cd*g338-384* wild-type) / (cd*g384* cd*384*)** | 208.5 | 289.56 |  | 3.84 | 0 |
| **(cd*g384_del* wild-type) / (cd*g384* wild-type)** | 0.16 | 0.14 |  | -2.13 | 0.03 |
| **(cd*g384_del* wild-type) / (cd*g384* cd*225*)** | 0.11 | 0.11 |  | -2.17 | 0.03 |
| **(cd*g384_del* wild-type) / (cd*g225* cd*384*)** | 2.87 | 3.47 |  | 0.87 | 0.38 |
| **(cd*g384_del* wild-type) / (cd*g338-384* cd*384*)** | 1.12 | 1.34 |  | 0.1 | 0.92 |
| **(cd*g384_del* wild-type) / (cd*g384* cd*384*)** | 25.92 | 30.79 |  | 2.74 | 0.01 |
| **(cd*g384* wild-type) / (cd*g384* cd*225*)** | 0.72 | 0.64 |  | -0.37 | 0.71 |
| **(cd*g384* wild-type) / (cd*g225* cd*384*)** | 18.33 | 20.52 |  | 2.6 | 0.01 |
| **(cd*g384* wild-type) / (cd*g338-384* cd*384*)** | 7.15 | 7.89 |  | 1.78 | 0.07 |
| **(cd*g384* wild-type) / (cd*g384* cd*384*)** | 165.25 | 181.37 |  | 4.65 | 0 |
| **(cd*g384* cd*225*) / (cd*g225* cd*384*)** | 25.58 | 31.43 |  | 2.64 | 0.01 |
| **(cd*g384* cd*225*) / (cd*g338-384* cd*384*)** | 9.99 | 12.11 |  | 1.9 | 0.06 |
| **(cd*g384* cd*225*) / (cd*g384* cd*384*)** | 230.68 | 278.78 |  | 4.5 | 0 |
| **(cd*g225* cd*384*) / (cd*g338-384* cd*384*)** | 0.39 | 0.54 |  | -0.68 | 0.5 |
| **(cd*g225* cd*384*) / (cd*g384* cd*384*)** | 9.02 | 12.44 |  | 1.59 | 0.11 |
| **(cd*g338-384* cd*384*) / (cd*g384* cd*384*)** | 23.1 | 31.59 |  | 2.3 | 0.02 |

**Table S3. The inheritance bias observed for *cd^338-384^* and *cd^384^* is due to homing and not meiotic drive.** Full model summaries for estimates of inheritance rates from AGG1928 in crosses with *cd^g384^* and *cd^g338-384^*. Log-odds and 95% confidence intervals, and significance values are taken from a binomial glmm with 'logit’ link using sgRNA line.

| Predictors | Odds Ratios | 95% CI | P-Value |
| --- | --- | --- | --- |
| **(Intercept)** | 0.98 | 0.89 – 1.08 | 0.751 |
| **gRNA type [cd*g384*]** | 0.95 | 0.84 – 1.08 | 0.434 |
| **Observations** | 32 | | |

| **Transgenic** | **Estimate** |
| --- | --- |
| ***Cd^338-384^*** | 0.496 (0.472-0.52) |
| ***Cd^384^*** | 0.83 (0.462-0.505) |

**Table S4. Proportion of reads with out-of-frame and in-frame indels in in the F1 trans-heterozygous males of the *zpg^3’Cas9^;cd^g338-384^* , the *zpg^3’Cas9^*;*cd^g384^* , and the *zpg^3’Cas9^*;*cd^g225^*.** Proportions are expressed in percentage of the reads with indels. Source data is provided as a Source Data file.

|  | Proportion of out-of-frame indels | Proportion of In-frame indels | Total number of reads with indels/total aligned reads |
| --- | --- | --- | --- |
| *zpg^3’Cas9^;cd^g338-384^* | 70.06% | 29.94% | 33819/60438 |
| *zpg^3’Cas9^*;*cd^g384^* | 78.57% | 21.43% | 6630/10470 |
| *zpg^3’Cas9^*;*cd^g225^* | 71.72% | 28.28% | 9541/36536 |
| SDA-500 | 72.29% | 27.71% | 83/66002 |

**Table S5. Proportion of reads lacking single to all sgRNAs of total modified reads and total reads in the F1 trans-heterozygous males of the *zpg^3’Cas9^;cd^g338-384^* , the *zpg^3’Cas9^*;*cd^g384^* , and the *zpg^3’Cas9^*;*cd^g225^*.** Source data is provided as a Source Data file.

|  | Proportion of modified reads | Proportion of total reads |
| --- | --- | --- |
| *zpg^3’Cas9^;cd^g338-384^* |  |  |
| Lacking 1 sgRNA | 44.2% | 26.5% |
| Lacking 2 sgRNAs | 25.6% | 15.4% |
| Lacking 3 sgRNAs | 15.6% | 9.4% |
| Lacking 4 sgRNAs | 14.2% | 8.5% |
| Total | 99.6% | 59.8% |
| *zpg^3’Cas9^*;*cd^g384^* |  |  |
| Lacking sgRNA384 | 100% | 68.2% |
| *zpg^3’Cas9^*;*cd^g225^* |  |  |
| Lacking sgRNA225 | 100% | 29.1% |

**Table S6. List of primers.**

| Primer name | Sequence | Purpose |
| --- | --- | --- |
| LA247 | CGATTGATGAGTCATTTGTT | PCR and Sequencing of *cd^g384_del^* |
| LA299 | GGTAACCGAGACAATGGAGAAGCAAGAG | PCR and Sequencing of *cd^g384_del^* |
| LA4330 | CGCAGCCGAAACGTGTTCAACATTC | PCR and Sequencing of *cd^g338-384^*, *cd^g384_del^*, *cd^g225^* |
| LA4331 | AATGCGTTGTATCAAATGACACGGCG | PCR and Sequencing of *cd^g384_del^*, *cd^g225^* |
| LA4332 | GTCTGTTTAAACTTCACCGTCTTTCTGGG | PCR and Sequencing of *cd^g225^* |
| LA4333 | AGTTAAACCCGGACTATGGTGATGGC | PCR and Sequencing *cd^384^*, *cd^g384_del^*, *cd^g225^* |
| LA4335 | CCGTGGCTTGATTAAGTTGCTGCC | Sequencing *cd^225^* |
| LA4336 | GTACAGCGAGTAAGGATTGAACAGCATCC | PCR of cd^384^, *cd^g225^* |
| LA4337 | CAGATAATGCTCGACAGCTTTGTGCC | PCR and Sequencing of *cd^g384_del^* |
| LA4338 | AGTACATACATTCTCAACCGAAGGCGC | PCR and Sequencing of *cd^g338-384^*, *cd^g384_del^* |
| LA4902 | CGTTATCAACTTGAAAAAGTGGC | PCR and Sequencing of *cd^g338-384^*, *cd^g384_del^* |
| LA5160 | GACCCAAGAAAAAGCGGAAGGTGG | PCR and Sequencing of *cd^g384_del^* |
| LA5986 | ATCAAGCTCCTCTAGATCCGGTGGATCTTAC | PCR and Sequencing of *cd^g384_del^* |
| LA6101 | CTGACCAAGGAGATGACCATGAAGTACCGCATG | PCR and Sequencing of *cd^g384_del^* |
| LA6731 | GGTAACCGAGACAATGGAGAAGCAAGAG | PCR and Sequencing of *cd^g225^* |
| LA6791 | CCAGTTCGGTTATGAGCCGT | PCR and Sequencing of *cd^g225^* |
| LA6792 | AATGACATCATCCACTGATCG | PCR and Sequencing of *cd^g338-384^* |
| LA7592 | CTCGGACGACGAGCGTATGG | PCR and Sequencing of *cd^g225^* |
| LA7953 | ATTGAAGGCCCGCGATAGACTC | PCR and Sequencing of *cd^g225^* |
| LA8538 | TAGCGGCCAAGGTGGGATTTGTGGAGGATCGTGCTAC | PCR and Amplicon Sequencing of *zpg^3’Cas9^;cd^g225^* |
| LA9133 | TCGACGGAAGCACGTGGCGCCGAAA | PCR and Amplicon Sequencing of *zpg^3’Cas9^;cd^g225^* |
| LA8682 | TTTCGGCGCCACGTGCTTCCGTCGA | PCR and Amplicon Sequencing of *zpg^3’Cas9^;cd^g338-384^*, *zpg^3’Cas9^;cd^g384^*, SDA-500 |
| LA8684 | GCAGCTGTCGACCGTCCGGTGTT | PCR and Amplicon Sequencing of *zpg^3’Cas9^;cd^g338-384^*, *zpg^3’Cas9^;cd^g384^*, SDA-500 |

**Table S7. Number of reads for each sample analysed by CRISPResso2.**

| Sample name | Reads in inputs | Reads after preprocessing | Reads aligned |
| --- | --- | --- | --- |
| *zpg^3’Cas9^;cd^g338-384^* | 66158 | 63256 | 60438 |
| SDA-500 | 71533 | 68186 | 66002 |
| *zpg^3’Cas9^*;*cd^g384^* | 45512 | 42529 | 10470 |
| *zpg^3’Cas9^*;*cd^g225^* | 47622 | 41612 | 36536 |
